# Invasive plants facilitated by socioeconomic change shelter vectors of scrub typhus and spotted fever

**DOI:** 10.1101/661892

**Authors:** Chen-Yu Wei, Jen-Kai Wang, Han-Chun Shih, Hsi-Chieh Wang, Chi-Chien Kuo

## Abstract

**Background:** Ecological determinants of most emerging vector-borne diseases are little studied, particularly for neglected tropical disease; meanwhile, although socioeconomic change can have significant downstream effect on human risks to vector-borne diseases via a change in land cover, particularly facilitating the invasion of exotic plants, related studies remain very scarce. Scrub typhus and spotted fever are neglected diseases emerging around the globe and are transmitted by chigger mites and ticks, respectively, with small mammals as the primary hosts of both vectors.

**Methodology/Principal findings:** We investigated how invasion of *Leucaena leucocephala* plant after extensive abandonment of farmlands driven by industrialization in Penghu Islands of Taiwan affected abundance of chiggers and ticks by trapping small mammals in three types of habitats (invasion site, agricultural field, human residence) every two months for a year. Invasion sites sheltered more chiggers and ticks than the other two habitats; moreover, both vectors maintained higher abundance in early winter and populations of chiggers were more stable across seasons in invasion sites, suggesting that the invasive sites could be a temporary refuge for both vectors and might help mitigate the negative influence of unfavorable climate. Infective rates of etiologic agents in chiggers and ticks were also higher in invasion sites. Top soil temperature and relative humility were similar across the three habitats, but invasion sites harbored more *Rattus losea* rat, on which infested chiggers and ticks were more well fed than those from the most commonly trapped species (*Suncus murinus* shrew), implicating that abundance of superior hosts instead of microclimate, might determine the abundance of both vectors.

**Conclusions/Significance:** This study highlights an important but largely neglected issue that socioeconomic change can have unexpected consequence for human health mediated particularly through invasive plants, which could become a hotspot for emerging infectious diseases but usually are very hard to be eradicated. In the future, a more holistic perspective that integrates socioeconomy, land use, exotic species, and human health should be considered to fully understand potential emergence of vector-borne diseases.

**Author summary:** Understanding how environmental factors, such as land use change, affect risks to vector-borne diseases helps control and prevent human diseases, but ecological preference of vectors of most neglected diseases remain little investigated. In this study, we found that vectors of scrub typhus (chigger mites) and spotted fever (hard ticks), two emerging neglected diseases, were much more abundant in sites invaded by exotic plants than the other major land cover types in a small island of Taiwan; moreover, populations of chigger mite in invasion sites were more stable across seasons, suggesting that plant invasion sites could be a refuge for disease vectors under unfavorable climate. Higher abundance of chigger mites and ticks was related to higher abundance of a superior rodent host instead of a difference in soil micro-climate. More significantly, these invasive plants are facilitated by extensive abandonment of farmlands driven by industrialization and rural to urban migration, thus demonstrating an important but largely neglected issue that socioeconomic change, when mediated through a change in land cover, can have unexpected downstream effect on emerging neglected tropical diseases.

## Introduction

Many vector-borne diseases are emerging around the globe, but importance of ecological factors in driving these emergence, such as climate change and land use change, remains largely unconfirmed [1, 2], particularly when concerning neglected tropical diseases. There is growing concern that plant invasion can have unexpected consequence for human health, including risks to vector-borne diseases [3]. Limited studies revealed that exotic plants can increase or sometimes decrease abundance of disease vectors. For example, there were more tick vectors of Lyme disease in Japanese barberry invasion sites than in areas dominated by native shrubs [4–7]. Likewise, ehrlichiosis-transmitting ticks were more abundant in sites occupied by invasive Amur honeysuckle than in sites free of it [3]. Invasive plants could also benefit some mosquito species [8–11]. By contrast, exotic plants can reduce the survival of some ticks [12] or been less preferred oviposition sites for mosquito vectors of La Crosse virus [13].

However, these comparative studies were typically implemented within a limited period of a year, without further investigating whether the extent or direction of differential vector survival or abundance in invaded versus non-invaded habitats might vary with seasons. For example, invasive plants might help vectors endure unfavorable weather or season by maintaining more stable climatic conditions under dense vegetative cover or by providing refuges for vertebrates that act as hosts for some disease vectors (such as ticks and some mite species); if this is true, eradicating invasive plants will become more pressing when these plants can ameliorate the negative effects of extreme weather conditions under further climatic change. Furthermore, to better predict human risks to vector-borne diseases after plant invasion, elucidating mechanisms enhancing or suppressing disease vector is essential, but related studies remain very scarce (but see [3,10,12]). Abundance of Acari disease vectors like ticks that alter life cycles on vertebrate hosts and in the soil can generally be determined by abiotic and biotic factors [3]: the former includes an alteration of soil surface microclimate after plant invasion that can affect survival of questing ticks [12]; invasive plants can also help aggregate ticks by providing food or cover for their vertebrate hosts [3].

Scrub typhus and spotted fever are neglected diseases that are emerging around the globe [14, 15]. Scrub typhus is an acute and potentially lethal febrile disease transmitted by chigger mites (Trombiculidae) infective of the rickettsia *Orientia tsutsugamushi* (OT) and has long been thought confined to Asia and northern Australia [16]. However, this disease has recently been identified in South America (Chile and Peru [17–19]) and Africa (Kenya and Djibouti [20–22]), and is also emerging in some endemic regions, such as China and Korea [23–27]. The life cycle of chigger mites include the egg, larva, nymph, and adult; only the larval stage (chiggers thereafter) is parasitical, feeding primarily on rodents and is the only stage transmitting OT to humans, while the nymph and adult are free living in the soil, predating on arthropods [28–30]. Chigger mites are the only reservoir of OT [14, 31], with extremely high efficiency of transstadial (from larva to nymph to adult) and transovarial (from adult to progeny) transmission occurring within chiggers [32–33]. Because chigger mites spend >99% of their life cycle time in the soils [34], other than rodents as the main food resource of parasitical chiggers, the soil temperature and moisture also determine the abundance and distribution of chigger mites [29, 30].

Likewise, spotted fever is emerging around the globe and is transmitted primarily by hard ticks (Ixodidae) infective of spotted fever group (SFG) rickettsiae (*Rickettsia* spp.) [35, 36]. Similar to chigger mites, life stages of hard ticks include eggs, larvae, nymphs, and adults; however, unlike chigger mites with larvae as the only parasitical stage, larval, nymphal, and adult hard ticks are all parasitical, requiring blood meal from vertebrates to molt or lay eggs. Ticks are reservoirs of SFG rickettsiae, which can be transmitted transstadially and transovarially within ticks, and all three parasitical stages are capable of vectoring SFG rickettsiae to humans [37]. Like chigger mites, hard ticks spend a great proportion of time on the ground (>90%, [38]), so their population is also affected by soil temperature and humidity [38–40].

Meanwhile, socioeconomic change can cause a change in land use and land cover, including invasion of exotic plants. Farming continues to dramatically transform earth landscapes [41], but global abandonment of agricultural fields has also increased considerably since the 1950s [42]. Abandonment usually occurs in remote, marginal grazing and agricultural lands, where soils are poor and fertility and profits are low [43–46], and is driven primarily by socio-economic factors that lead to depopulation of rural areas [47], including such as industrialization, rural-urban migration, and urbanization [43-46,48,49]. These abandoned old fields, particularly degraded lands with strong cultivation legacy, are typically dominated by prolific and better competitive invasive plants that can impede the recovery of native plants [42,45,48,50]

The Penghu Islands, previously known as the Pescadores Islands, are located in the Taiwan Strait (Fig 1) and is comprised of 90 subtropical and tropical islands, with the largest island covering an area of 65 km^2^. The climate in Penghu is characterized by hot summer and dry and windy winter [51]; farmlands are usually surrounded by walls made of coral stones to fend off strong winds (Fig 2a). Moreover, due to the small size of islands, the lands are continuously rained with salty sea water, particularly during the windy winter; this had led to high soil salinity. Unfavorable climate and poor soil fertility immensely limit agricultural productivity. Industrialization of Taiwan beginning in the 1970s was accompanied in Penghu by a steep decline in the area of cultivated lands and proportion of workers that are farmers (Fig 3, adapted from [51]). The outcome is that most agricultural fields in Penghu are left abandoned. For example, as of 2016, about 70% of workers were in the service sector [52], and in 2015, 75% of farmlands was abandoned, the highest among all counties in Taiwan, far higher than the next highest county (around 30% abandonment rate [53]). These abandoned fields are invaded almost exclusively by a nitrogen-fixing legume, the exotic white popinac *Leucaena leucocephala*, which is among 100 of the world’s worst invasive alien species listed by IUCN Invasive Species Specialist Group [54]. *L. leucocephala*, native to Central America, is introduced worldwide as firewood or fodder plants and can become highly invasive in disturbed sites with dry and poor soils; they can prevent native vegetation recovery by forming dense thickets that are also very difficult to be eradicated (Global Invasive Species Database, IUCN, http://www.iucngisd.org/, accessed October 17, 2018). In Penghu, the density of *L. leucocephala* can reach 30,000 to 50,000 stands per hectare [55] and eradicating *L. leucocephala* has been a priority for local government.

**Fig. 1.**
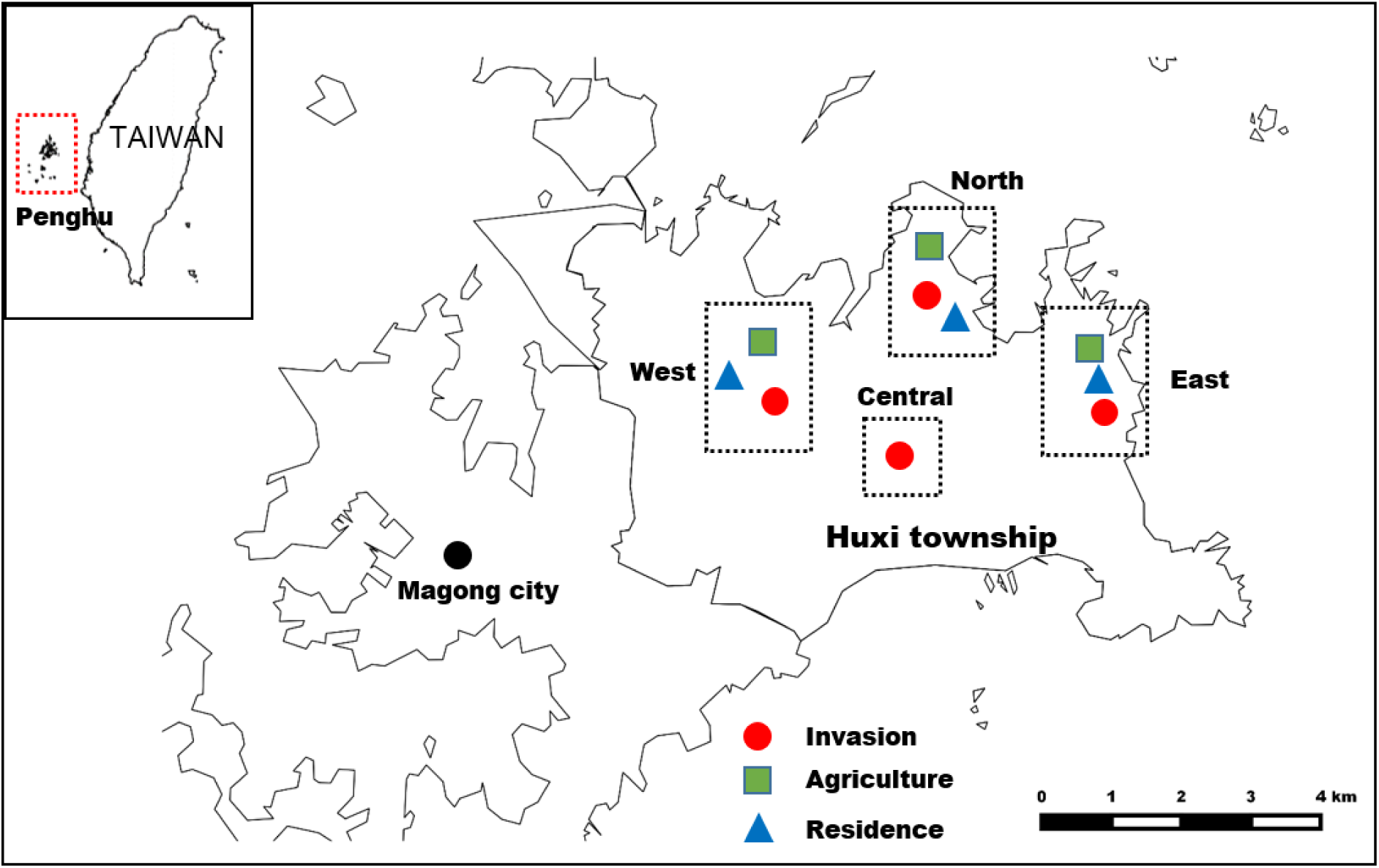
Study sites in Huxi township of Penghu Islands. The maps were created by the authors with QGIS 2.12.2-Lyon by QGIS Development Team.

**Fig. 2.**
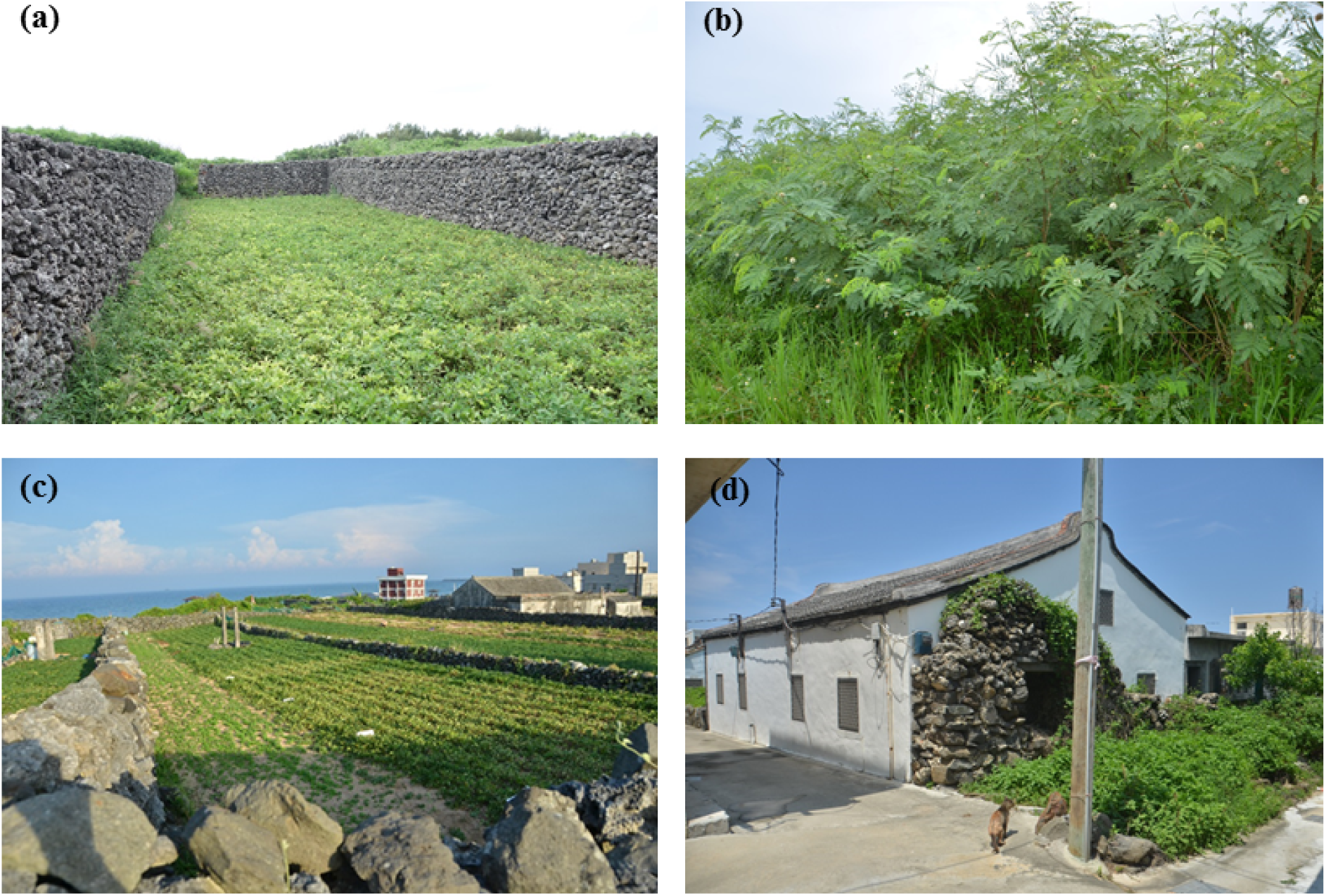
Habitats in Penghu Islands. (a) Farmers typically use coral stones to build walls for fending off strong wind during the winter; (b) *Leucaena leucocephala* invasion sites; (c) agricultural fields; (d) human residence sites.

**Fig. 3.**
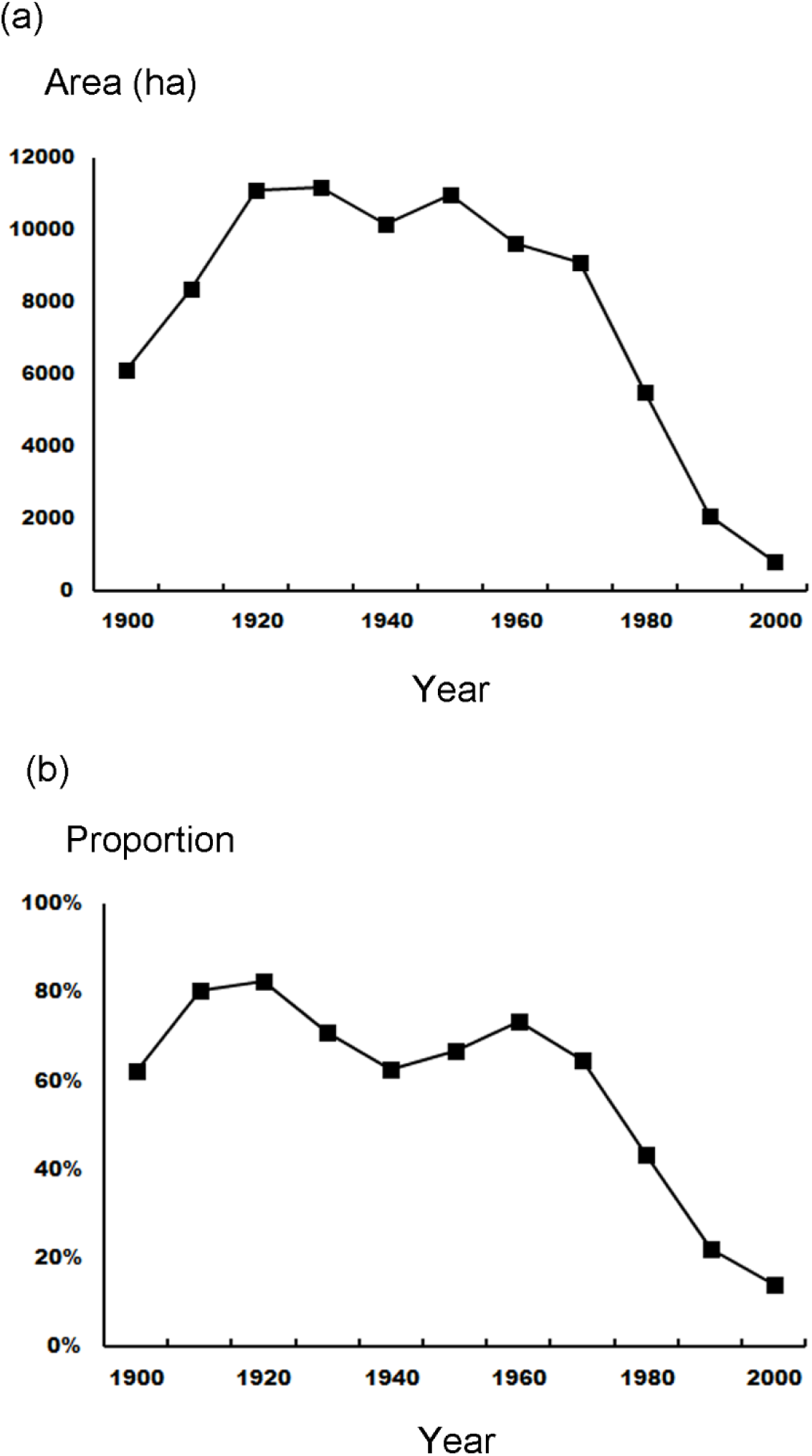
(a) The area of cultivated lands (ha) and (b) proportion of workers that were farmers in Penghu Islands.

Penghu Islands, at the same time, is the hotspot of scrub typhus, with the highest number of human cases of notifiable scrub typhus among all counties in Taiwan for the past ten years (2008-2017, Taiwan Centers for Disease Control, https://nidss.cdc.gov.tw/, accessed October 17, 2018). Despite that scrub typhus in Penghu was intensively studied by the U.S. Naval Medical Research Unit Two in the 1960s and 1970s [56–62], these studies have never focused on habitat difference in chigger vectors, including evaluating the significance of invasive plants. Likewise, SFG rickettsiae have been detected in hard ticks and small mammals in Penghu, but these were based on very limited sample size (3 and 30 for ticks and small mammals, respectively) [63–64] and without investigating the differential importance of habitats in sheltering disease vectors.

Although socioeconomic change can have significant downstream effect on human risks to vector-borne diseases via a change in land use and vegetative community, related studies remain very limited. Here, we investigated (1) whether the invasion of *L. leucocephala*, facilitated by socioeconomic change, creates ideal habitats for chiggers and hard ticks. Comparison among habitats was also implemented across seasons to assess if invasive plants help sustain vector populations under unfavorable climate. (2) Furthermore, the importance of abiotic versus biotic factors in determining differential vector populations among habitats was evaluated. (3) Lastly, because feeding success of chiggers and hard ticks could vary with host species identity [65, 66], we also investigated whether vectors attracted by some host species have low feeding success so that preserving these host species might lower vector population and disease risks to human.

## Materials and methods

### Ethical statement

All animal handling procedures were approved by the National Taiwan Normal University Animal Care and Use Advisory Committee (permit number NTNU-104016), which adheres to Guideline for the Care and Use of Laboratory Animals established by Council of Agriculture, Taiwan.

### Study sites, small mammal trapping, and ectoparasite collection

This study was implemented in the Huxi Township (Fig 1), where more than half of all Penghu scrub typhus human cases occurred in the last ten years (2008-2017, Taiwan Centers for Disease Control, https://nidss.cdc.gov.tw/, accessed October 17, 2018). A total of ten study sites were investigated, including four *L. leucocephala* invasion sites, three agricultural fields, and three human residence sites (traps placed outdoor instead of indoor) (Fig 2b-d) in four different parts (east, west, north, and central) of Huxi (Fig 1). From December 2016 to October 2017, small mammal traps were set up in each of these ten sites every two months. In December 2016 and February 2017, 30 Sherman small mammal traps (26.5 × 10.0 × 8.5 cm) were deployed in each site, while starting from April 2017, each site was supplemented with three meshed traps (27.0 × 16.0 × 13.0 cm) in addition to the 30 Sherman traps to increase sampling efforts. During each capture session, traps were baited with sweet potatoes covered with peanut butter and were open for three consecutive nights.

Trapped small mammals, including shrews and rodents, were identified to species, sexed, measured for body weight, body length, and tail length, and examined for ectoparasite infestation. Skins with attached chiggers were removed from host animals with tweezers and placed in vials; 100% ethanol were added after 2-3 days when chiggers have released themselves from the skins to preserve intact oral parts for later species identification. Ticks were carefully collected with tweezers and preserved in 100% ethanol. All infested chiggers and ticks were collected and stored in −20°C refrigerator for subsequent molecular determination. Large rodent species, including *Rattus losea* and *Rattus norvegicus*, were each implanted with a radio-frequency identification chip (Watron Technology Corporation, Hsinchu, Taiwan) for individual identification before release. Smaller species, including *Suncus murinus* and *Mus musculus*, were unable to be permanently marked without difficulty, so were released without being marked.

### Species identification and engorgement degree of chiggers and ticks

Chiggers were slide-mounted in Berlese fluid (Asco Laboratories Ltd, Manchester, U.K.) and morphologically identified to species under a compound microscope following [67]. Ticks were morphologically identified to species and life stages (larva, nymph, male adult, female adult) under a dissecting microscope following published keys [68] and when species unrecognized, molecularly identified by comparing 12S rDNA and 16S rDNA to known species following [69, 70]. All ticks were identified, while due to the very large number of chiggers, only a portion of chiggers (>25%) from each individual host were examined.

Degree of engorgement of chiggers and ticks was compared among host species. Engorgement degree of a chigger was represented by the increase in idiosoma area relative to the one with the smallest idiosome area, which was calculated based on the ellipse equation [66]. Engorgement degree was firstly averaged within each host individual before subsequent interspecific comparison. Engorgement degree of ticks was divided into three categories: non-engorged, half-engorged, and fully engorged. Unlike chiggers, interspecific comparison of ticks was based on individual tick irrespective of whether collected from the same host individual.

### Detection of OT in chiggers and *Rickettsia* in ticks

Due to the minute size of chiggers, a sum of 30 chiggers from the same host individual was pooled for detection of OT with nested polymerase chain reaction (PCR) following [71], which targeted the well conserved 56-kDa type specific antigen located on the OT outer membrane. Laboratory OT strain and phosphate-buffered saline (PBS) solution were used as positive and negative controls, respectively. Tick was individually assayed for *Rickettsia* infection with nested PCR following [64], which targeted the 120- to 135-kDa surface antigen (*ompB*) and citrate synthase (*gltA*). Laboratory *R. rickettsii* antigen and PBS solution were used as positive and negative controls, respectively.

### Top soil temperature and relative humidity

Temperature and relative humidity of the top soil were recorded from December 2016 to October 2017 by placing a data logger (WatchDog, Spectrum Technologies Inc., East Plainfield, Illinois) on the ground in each of the 10 study sites. Measurements were recorded at an interval of 30 minutes.

### Statistical Analyses

Because *S. murinus* and *M. musculus* were not individually marked, only results from the 1^st^ day of capture in each three-day trapping session were used to calculate capture rate (unique individuals/trap-nights) and ectoparasite load (number of all ectoparasites/number of all host individuals); including the 2^nd^ and 3^rd^ day capturing results might overestimate capture success (when host individuals were recaptured) and underestimate ectoparasite load (when recaptured hosts had been removed of chiggers and ticks the previous day/days). However, when tallying the total number of ectoparasites in each site, results from all three days were included; including even recaptured host individuals will not or will only slightly increase total ectoparasite numbers (as recaptured hosts had been removed of chiggers and ticks the previous one or two days). By contrast, because *R. losea* and *R. norvegicus* was individually marked, results from all three days were included in the analyses (but excluding recaptured individuals in the same 3-night trapping session).

Loads of chiggers and ticks were compared among host species with negative binomial generalized linear model to account for overdispersion of data, and significant difference was evaluated based on the 95% Wald confidence interval. When comparing engorgement degree of chiggers among host species, normality and homogeneity of variance were confirmed with Shapiro– Wilk and Levene tests, respectively. Data were transformed when necessary and if homogeneity of variance cannot be fulfilled even after transformation, Welch’s ANOVA was implemented followed by Games-Howell post hoc test. When comparing engorgement degree of ticks among host species, as well as whether host species vary in their relative importance among habitats in hosting ticks, Fisher-Freeman-Halton’s test with 100,000 Monte Carlo permutations were implemented, and if significant, followed by pair-wise Fisher-Freeman-Halton’s test with Bonferroni correction. When investigating whether host species vary in their relative importance among habitats in hosting chiggers, and whether host species composition among habitats, Pearson chi-square test was applied.

We investigated difference in chigger and tick abundance, as well as variation in *R. losea* capture rate among regions, habitats, and months with generalized estimating equations (GEE) using negative binomial log link function, with site as the subject, and each bi-monthly sampling as repeated measures within the site (ten sites, each with six sampling, so overall 60 samples). Region, habitat, month, and habitat*month were the fixed factors, and significance of difference was determined based on 95% Wald confidence interval of estimated marginal means. The structure of the correlation matrix was selected based on the lowest quasi-likelihood under independence model criterion value. To avoid that the Hessian matrix is not positively definite so that reliable results can’t be attained, the dependent variable (chigger, tick, or *R. losea* capture rate) was set as 0.001 when the original value was zero. GGE model was also implemented when comparing *S. murinus* capture rate among explanatory variables except that a normal distribution function was instead applied. We also calculated the coefficient of variation (CV=standard deviation divided by mean) for chigger and tick abundance across months for each habitat type, represented by the mean of ectoparasite abundance in each site of the same habitat. For example, the abundance of chiggers in *L. leucocephala* invasion sites in December 2016 was represented by the average of the chigger abundance of the four *L. leucocephala* invasion study sites surveyed in that month.

When comparing prevalence of infection of OT and *Rickettsia* among host species and habitats, the more robust bootstrapped logistic regression was applied [72] instead of the conventional logistic regression that could cause biased results when sample size is small [73]; 95% confidence interval was estimated with 10,000 permutations. Data are given as the mean ± 1 standard error (SE). All the procedures were implemented in SPSS Statistics version 19.0 (IBM Corp.).

## Results

### Small mammal composition and chigger and tick infestations

A total of 1,345 small mammals of four species were captured out of a sampling effort of 5,760 trap-nights. The *S. murinus* shrew was the most abundant (capture rate = 0.198 individuals/trap-nights), followed by the rodents *M. musculus* (0.071), *R. losea* (0.029), and *R. norvegicus* (0.001).

A sum of 42,198 chiggers were collected, primarily from *R. losea* (76.3% of total), and to a less extent from *S. murinus* (18.7%), *M. musculus* (2.9%), and *R. norvegicus* (2.1%). Overall, chigger load (number of chiggers/number of host individuals) was significantly higher on *R. losea* (192.9±20.2 chiggers, mean ± 1SE, n=167) and *R. norvegicus* (110.1±45.4, n=8) than on *S. murinus* (9.6±2.0, n=380) and *M. musculus* (2.8±1.2, n=122) (negative binomial generalized linear model, all *p* < 0.05) (Fig 4a).

**Fig. 4.**
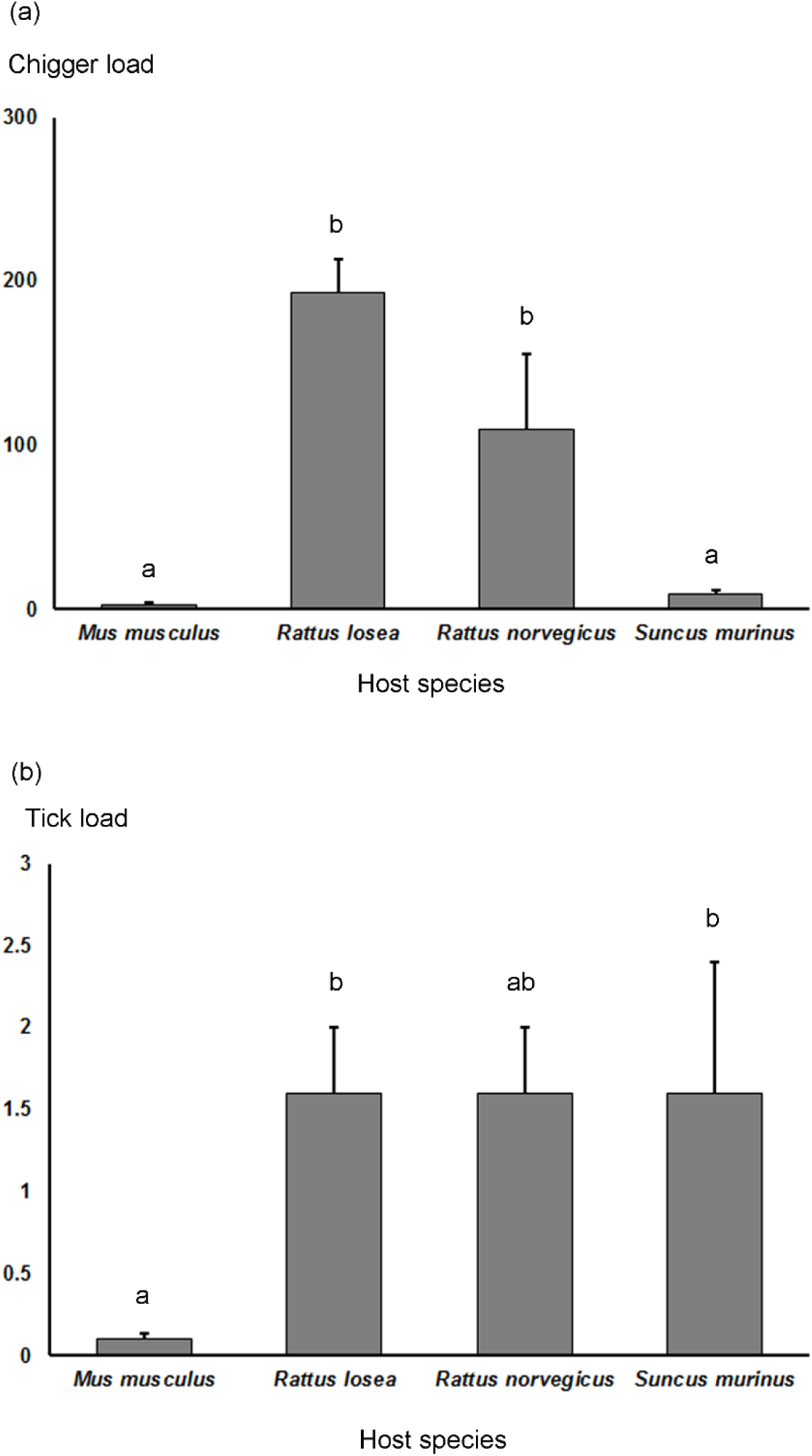
Mean load of (a) chiggers and (b) ticks on different host species in Penghu Islands from December 2016 to October 2017. Different letters represent significant difference. Error bar +1SE.

A total of 1,049 ticks were collected, including 486 larvae (46.3%), 288 nymphs (27.5%), and 275 adults (26.2%). These were mostly collected from *S. murinus* (69.0% of total), followed by *R. losea* (25.5%), *M. musculus* (4.8%), and *R. norvegicus* (0.7%). Tick load was significantly higher on *R. losea* (1.6±0.4, n=167) and *S. murinus* (1.6±0.8, n=380) than on *M. musculus* (0.1±0.04, n=122) (both *p* < 0.05), while *R. norvegicus* (1.6±0.4, n=8) was similar as the other three species (all *p* > 0.05) (Fig 4b).

### Species identification and engorgement degree of chiggers and ticks

A total of 10,815 chiggers were slide-mounted for species identification, including 1,035 chiggers (9.6% of total) that can’t be reliably identified due to specimens inadequately prepared. The other 9,780 successfully identified chiggers, including 324 chiggers from 25 *M. musculus*, 7,535 chiggers from 114 *R. losea*, 209 chiggers from six *R. norvegicus*, and 1,712 chiggers from 71 *S. murinus* were all *Leptotrombidium deliense*. Engorgement degree varied among host species (Welch’s ANOVA, F_3, 20.8_=164.1, *p* < .001), with chiggers on *R. losea* (10.4±0.2 × 10^4^ µm^2^) and *R. norvegicus* (11.3±1.4 × 10^4^ µm^2^) more engorged than those on *M. musculus* (4.9±0.5 × 10^4^ µm^2^) and *S. murinus* (3.9±0.2 × 10^4^ µm^2^) (Games-Howell test, all *p* < 0.005) (Fig 5a).

**Fig. 5.**
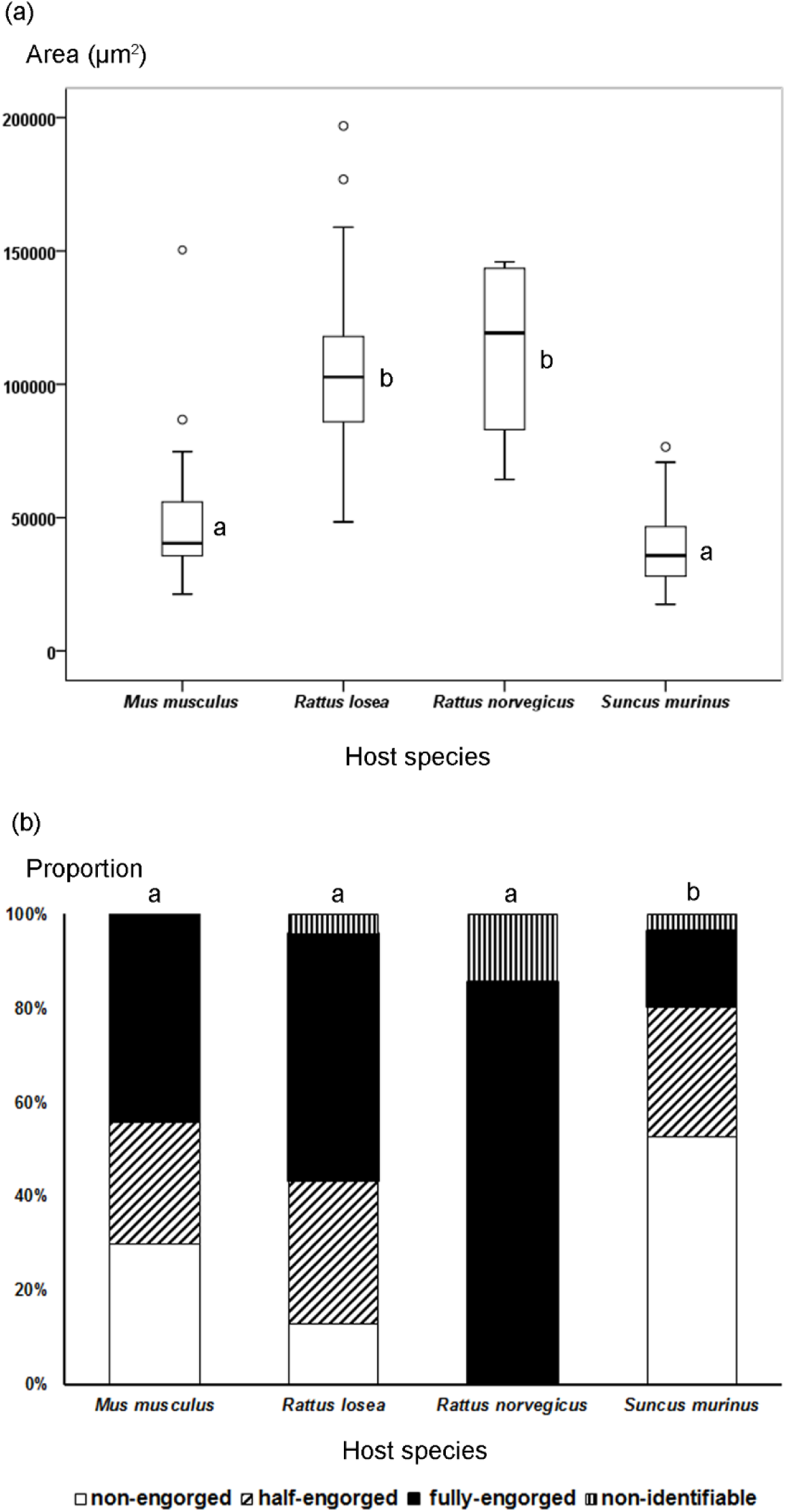
Engorgement degree of (a) chiggers and (b) ticks collected from different host species in Penghu Islands from December 2016 to October 2017. Different letters represent significant difference.

The 1,049 collected ticks were comprised predominantly (99.1%) of *Ixodes granulatus*; the other small proportion (0.9%) was *Amblyomma testudinarium*. More than half of the ticks (52.6%) collected from *S. murinus* were non-engorged while only 16.3% were fully engorged; on the contrary, 52.6% of ticks collected from *R. losea* were fully engorged and only 13.0% were non-engorged (Fig 5b). Engorgement degree varied among the four host species (Fisher-Freeman-Halton’s test, *p* < 0.001), with *S. murinus* differed from the other three species (all *p* < 0.05, after Bonferroni correction), while there was no difference among *R. losea*, *M. musculus*, and *R. norvegicus* (all *p* > 0.05) (Fig 5b).

### Variation in chigger abundance across regions, habitats, and months

The sum of chiggers collected from all mammal hosts or uniquely from *R. losea* both varied among regions, habitats, and months (GEE, all *p* < 0.001), and there was an interaction between habitat and month (both *p* < 0.001). There were more chiggers in the eastern region than the other parts of the study area (all *p* < 0.05) (Figs 6a-b). Invasion sites sheltered more chiggers than the other two habitats in most months, significantly in December (all *p* < 0.05) (Figs 7a-b). The coefficient of variation (CV) in chigger abundance across months was both lower in invasion sites (CV=0.79, 0.79; all mammals, only from *R. losea*, respectively) than in agricultural fields (1.07, 1.01) and residence sites (1.56, 1.61).

On the other hand, number of chiggers collected solely from *S. murinus* differed among regions and months (both *p* < 0.001) but not among habitats (*p* > 0.05), and there was an interaction between habitat and month (*p* < 0.001). There were significantly more chiggers in the eastern region than the north and west regions (both *p* < 0.05) but not the central (*p* > 0.05) (Fig 6c). There was no significant differences among the three habitats within the same month (all *p* > 0.05) (Fig 7c), although human residence sites in June sheltered significantly more chiggers than the other two habitats in most other months (but not in June). The CV value was lower in agricultural fields (1.35) than in invasion sites (CV=1.61) and human residence sites (1.93).

**Fig. 6.**
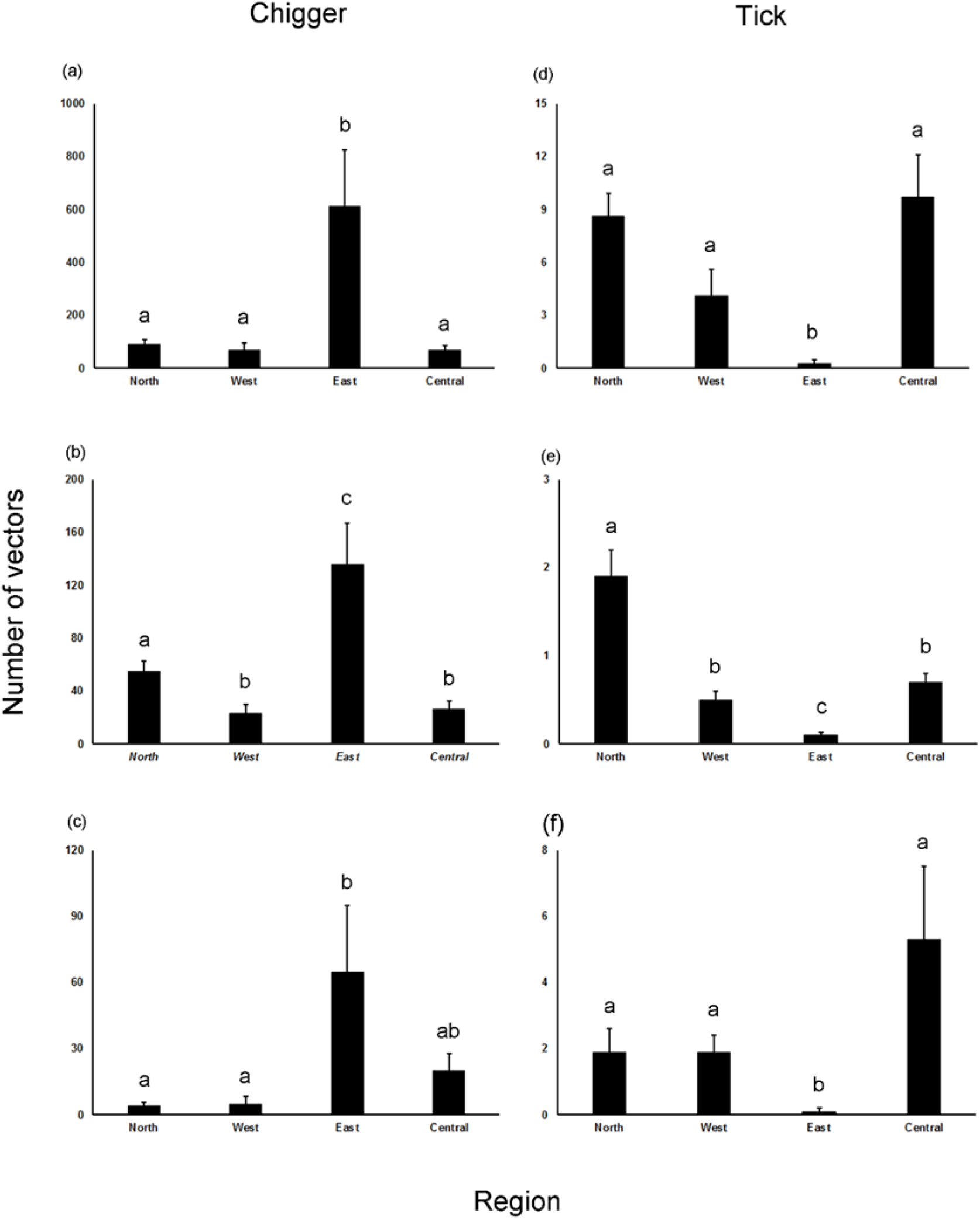
Number of vectors collected from mammal hosts per study site in different regions of Penghu Islands from December 2016 to October 2017. Chiggers collected from (a) all mammals combined; (b) *Rattus losea*; (c) *Suncus murinus*. Ticks collected from (d) all mammals combined; (e) *Rattus losea*; (f) *Suncus murinus*. Different letters represent significant difference. Error bar +1SE.

**Fig. 7.**
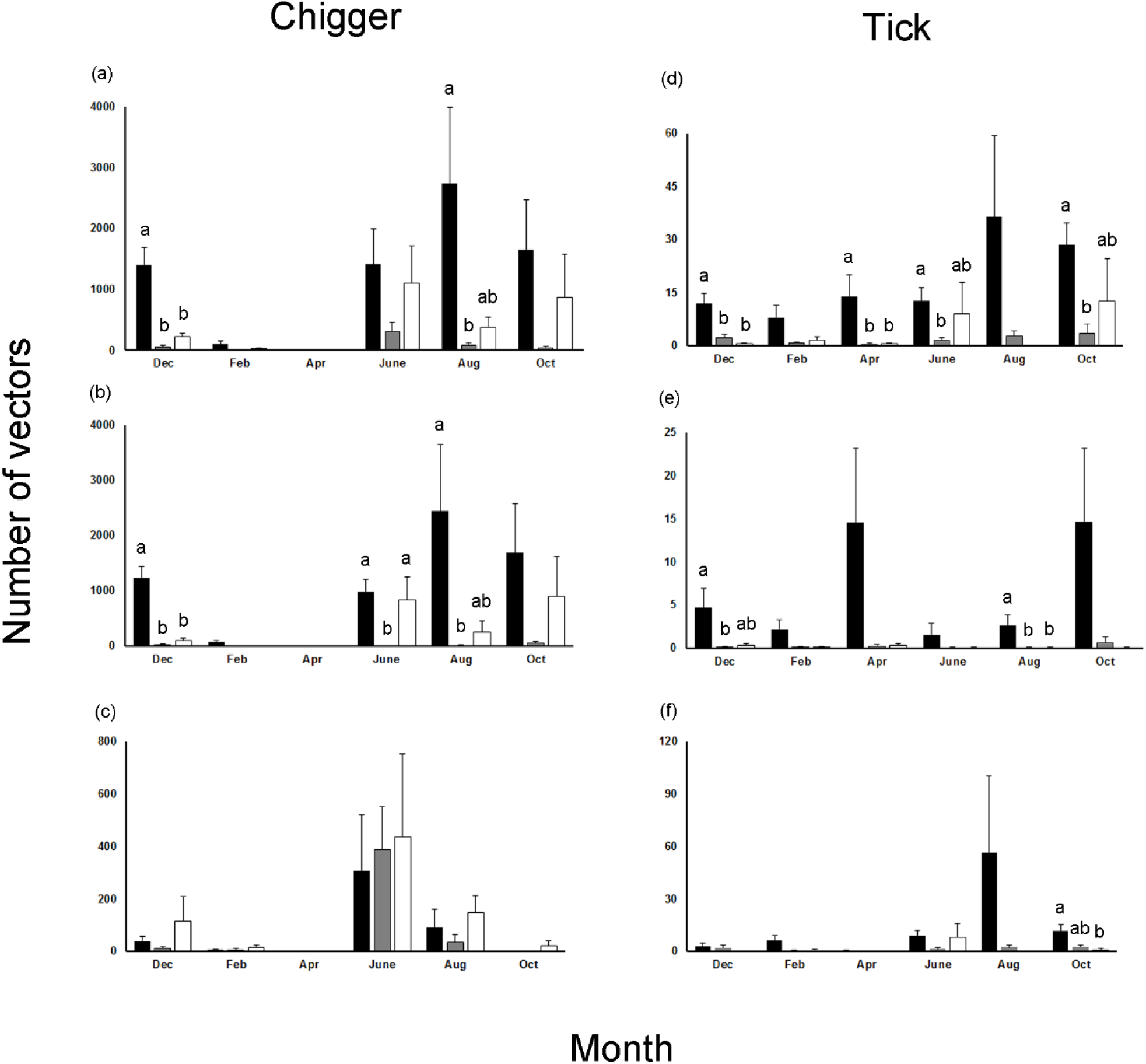
Number of vectors collected from mammal hosts per study site in different habitats and months in Penghu Islands from December 2016 to October 2017. Chiggers collected from (a) all mammals combined; (b) *Rattus losea*; (c) *Suncus murinus*. Ticks collected from (d) all mammals combined; (e) *Rattus losea*; (f) *Suncus murinus*. Different letters represent significant difference; statistical comparisons among habitats implemented only within months and letters were denoted only when significant difference was found. Black: *Leucaena leucocephala* invasion site; grey: human residence site; white: agricultural fields.

Within the *L. leucocephala* invasion sites and agricultural fields, chiggers were collected primarily from *R. losea* (89.4%, 71.2%; respectively), whereas in the human residence sites, chiggers were chiefly retrieved from *S. murinus* (84.1%); hosts vary in their relative contribution to feeding the chiggers among the three habitats (chi-square test, *χ*^2^= 16835.8, *p* < 0.001).

### Variation in tick abundance across regions, habitats, and months

Ticks collected from all mammal hosts, solely from *R. losea*, or exclusively from *S. murinus* all varied among regions, habitats, and months (GEE, all *p* < 0.001), and there was an interaction between habitat and month (*p* < 0.001). In most cases, the eastern region sheltered significantly fewer ticks than the other regions (all *p* < 0.05) (Figs 6d-f). Invasion sites sheltered significantly more ticks than the other two habitats in December and April and in August for all mammals (all *p* < 0.05, Fig 7d) and for *R. losea* only (all *p* < 0.05, Fig 7e), respectively, but mostly without significant difference for ticks collected solely from *S. murinus* (Fig 7f). The CV value for tick abundance across months was lower in agricultural fields (CV=0.58) than in residence (0.62) and invasion sites (0.80) for all mammals, lower in invasion sites (0.75) than in agricultural fields (1.18) and residence sites (1.03) for *R. losea* only, and lower in residence sites (CV=0.71) than in invasion sites (1.22) and agricultural fields (1.40) for *S. murinus* only.

Within the *L. leucocephala* invasion and human residence sites, ticks were collected principally from *S. murinus* (69.5%, 73.6%; respectively), while in the agricultural fields, ticks were collected equally from *R. losea* and *S. murinus* (both 40%); hosts vary in their relative importance among the three habitats (Fisher-Freeman-Halton’s test, *p* < 0.001).

### Variation in *R. losea* and *S. murinus* capture rate across regions, habitats, and months

Capture rate of *R. losea* varied among habitats and months (GEE, both *p* < 0.001), but not regions (*p* > 0.05), and there was an interaction between habitat and month (*p* < 0.001). Invasion sites sheltered more *R. losea* than the other two habitats for each month, significantly in December, February, and October (Fig 8a). On the other hand, capture rate of *S. murinus* varied among regions, habitats, and months (all *p* < 0.001), and there was an interaction between habitat and month (*p* < 0.001). Both eastern and central regions harbored more *S. murinus* than western and north regions (all *p* < 0.05). Human residence sites sheltered more *S. murinus* than the other two habitats for each month, significantly in June and August (Fig 8b).

**Fig. 8.**
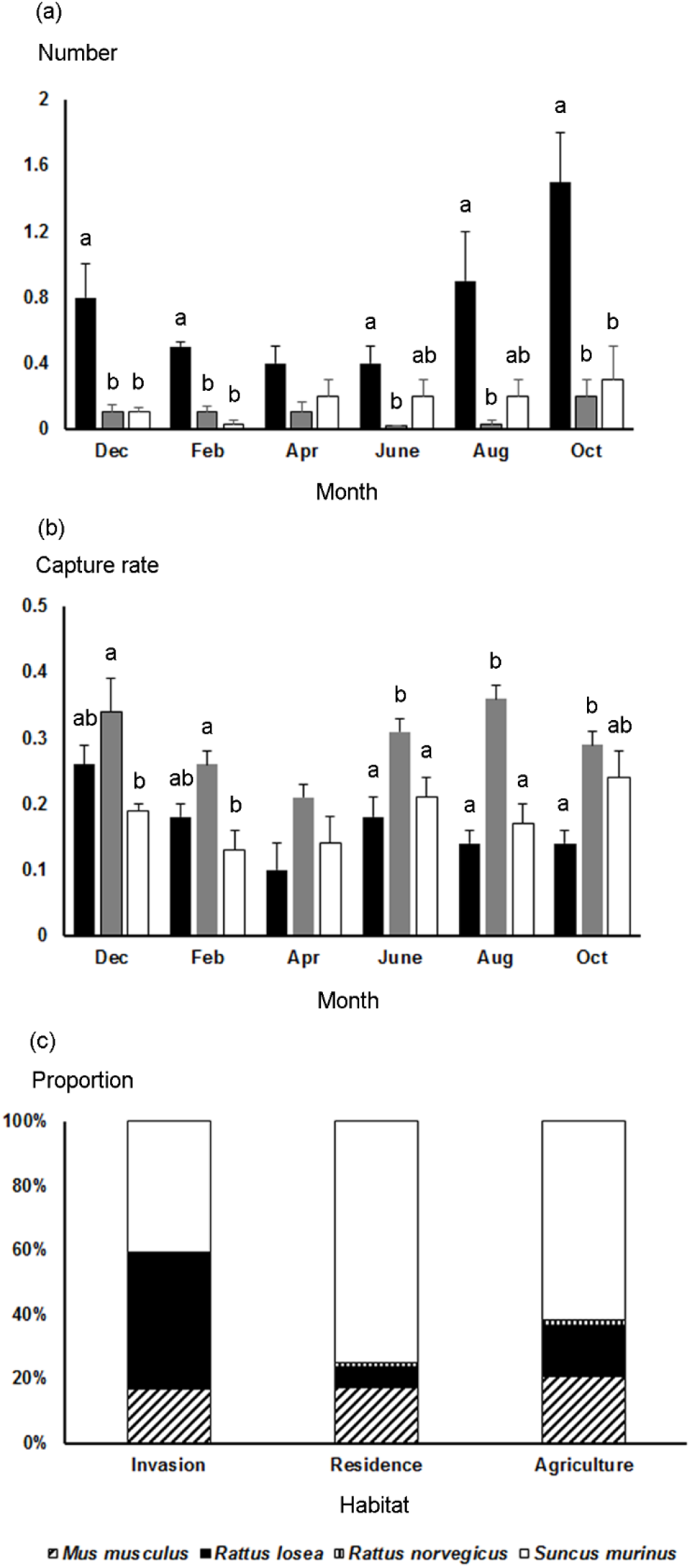
Abundance of small mammals in different habitats in Penghu Islands from December 2016 to October 2017. (a) Number of *Rattus losea*; (b) capture rate of *Suncus murinus*; (c) small mammal species composition. Different letters represent significant difference. For (a), (b), statistical comparisons among habitats implemented only within sampled months and letters were denoted only when significant difference was found. Black: *Leucaena leucocephala* invasion site; grey: human residence site; white: agricultural fields.

There were more small mammal captures in invasion (mean=77.5 individuals per site) and human residence sites (71) than in agricultural fields (51). In human residence sites, small mammals were comprised mainly of *S. murinus* (74.8% of total captures), followed by *M. musculus* (17.3%). The pattern was similar in the agricultural fields where *S. murinus* and *M. musculus* accounted for 61.4% and 20.9% of total captures, respectively. In comparison, in invasion sites, *R. losea* and *S. murinus* were the most common species, comprising respectively 41.6% and 40.6% of total captures (Fig 8c). Species composition varied among habitats (*χ*^2^= 99.4, *p* < 0.001).

### Prevalence of OT in chiggers and *Rickettsia* in ticks

A total of 154 pools of *L. deliense* chiggers were assayed for OT infections, with an overall prevalence of 21.4% (33/154). Prevalence was higher when chiggers were collected from *R. losea* (28.6%, 30/105) than from *S. murinus* (5.1%, 2/39) and *M. musculus* (0%, 0/7) (bootstrapped logistic regression, both *p* < 0.05), whereas chiggers from *R. norvegicus* (33.3%, 1/3) were the same as those from *R. losea* and *S. murinus* (both *p* > 0.05), but higher than *M. musculus* (*p* < 0.05). For chiggers on *R. losea*, prevalence of OT infection was higher when *R. losea* was trapped in *L. leucocephala* invasion (34.1%, 28/82) and human residence sites (28.6%, 2/7) than in agricultural fields (0%, 0/16) (both *p* < 0.05), but was similar between the first two habitats (*p* > 0.05). For chiggers on *S. murinus*, the prevalence was higher in invasion sites (14.3%, 1/7) and agricultural fields (7.7%, 1/13) than in human residence sites (0%, 0/19) (both *p* < 0.05), while there was no difference between the first two habitats (*p* > 0.05).

A sum of 180 *I. granulatus* ticks was individually tested for *Rickettsia* infections, with an overall prevalence of 8.3% (15/180). Prevalence was higher when ticks were collected from *M. musculus* (6.5%, 2/31), *R. losea* (12.3%, 7/57), and *S. murinus* (6.7%, 6/89) than from *R. norvegicus* (0%, 0/3) (all *p* < 0.05); there was no differences among the first three host species. Within *R. losea*, prevalence of *Rickettsia* infection in ticks was higher when *R. losea* was trapped in *L. leucocephala* invasion sites (15.6%, 7/45) than in agricultural fields (0%, 0/6), and human residence sites (0%, 0/6) (both *p* < 0.05); for ticks on *S. murinus*, prevalence was higher in invasion (7.8%, 5/64) and human residence sites (5.9%, 1/17) than in agricultural fields (0%, 0/8) (both *p* < 0.05), while there was no difference between the first two habitats.

### Top soil temperature and relative humidity

There was no significant difference in the monthly mean, minimum, and maximum temperatures among the three habitats for each of the 11 months (Fig 9a, ANOVA, all *p* > 0.05). There was also no difference among habitats (all *p* > 0.05) in the variation in monthly temperature (monthly maximum minus minimum) except in April when invasion sites had higher variation than the agricultural fields (*p* < 0.05) (Fig 9b). In terms of relative humidity, there was no significant difference in monthly mean, minimum, maximum humidity, and variation in humidity among the three habitats for each of the 11 months (Figs 9c-d, ANOVA, all *p* > 0.05).

**Fig. 9.**
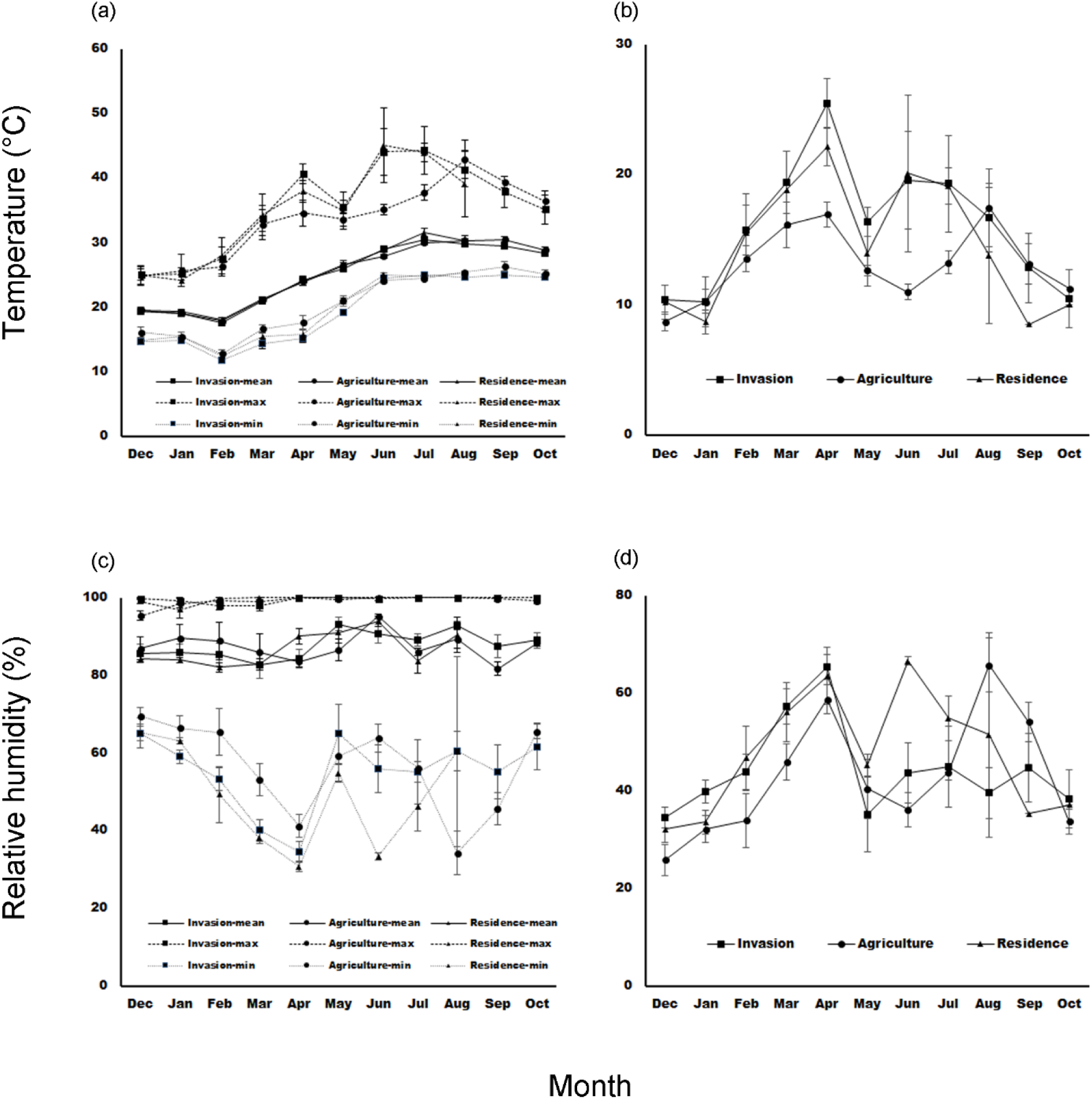
Monthly variation in temperature and relative humidity in different habitats in Penghu Islands from December 2016 to October 2017. (a) Minimum, mean, and maximum temperature; (b) fluctuation in monthly temperature (monthly maximum minus minimum); (c) minimum, mean, and maximum relative humidity; (d) fluctuation in monthly relative humidity (monthly maximum minus minimum).

## Discussion

More chiggers and ticks were collected from *L. leucocephala* invasion sites than from agricultural fields and human residence areas (Fig 7a, d); in addition, prevalence of infection of OT in chiggers and *Rickettsia* in ticks was higher when vectors were collected from invasion sites than the other two habitats, demonstrating that the proliferation of invasive *L. leucocephala*, encouraged by abandonment of marginal agricultural fields after industrialization in Taiwan, has created hotspots for scrub typhus and spotted fever. Moreover, *L. leucocephala* invasion sites maintained significantly higher number of disease vectors during early winter (December) than the other two habitats (Fig 7a, d). We also found that the *R. losea* rodent and the *S. murinus* shrew were the primary hosts of chiggers and ticks, but feeding success of both vectors was much higher on *R. losea* than on *S. murinus* (Fig 5), indicating that *R. losea* is a more ideal host than *S. murinus*. Lastly, there was little difference in top soil temperature and moisture among the three habitats (Fig 9), but more *R. losea* in invasion sites (Fig 8a), suggesting that higher abundance of chiggers and ticks in invasion sites might be partially related to residence of larger number of the ideal rodent host.

In this study, more than three quarters of chiggers were collected from *R. losea*, and degree of engorgement in *R. losea* was 2.7-fold that in *S. murinus*, indicating that *R. losea* is the most important host of chiggers in Penghu Islands, which is in agreement with our previous large scale study that *R. losea* is the primary host of chiggers across Taiwan [74]. In addition, in contrast to other pathogens, such as *Borrelia burgdorferi*, in which transovarial transmission rarely occurs and vertebrate hosts are necessary for re-infecting vectors [75], transovarial transmission of OT is high in chigger mites [33] and chigger mites are the only reservoir of OT [14], so species identity of hosts will not determine infectivity of feeding chiggers (i.e. relative to *R. losea*, *S. murinus* will not be more critical in maintaining OT circulation).

Although *L. deliense*, which is the primary chigger species vectoring OT in Southeast Asia [30], was also recognized as the dominant species in Penghu in this study as previous studies have shown [57, 62], past studies have instead identified *S. murinus* as the primary host of chiggers [56, 57]. The reason for such inconsistency is unclear, particularly when both previous studies did not document in which habitat type the traps were set up. One possibility is that *L. leucocephala* was not widespread in the 1960s so there were much fewer *R. losea* at that time. On the other hand, our finding that *S. murinus* hosted >80% of chiggers specifically in human dwelling areas is similar to the result of [57] that 70% of chiggers were collected from *S. murinus*, suggesting that past studies might limit the survey to human residence areas. This current study, after including other habitat types, has instead revealed *R. losea* as the most critical host of chiggers in Penghu. It should be noted, nevertheless, that more human activities surrounding human residence area than *L. leucocephala* invasion sites suggests that chiggers active in human residence area may be more pivotal in determining human risks to scrub typhus. In Penghu, *L. leucocephala* is so widespread that villages are typically surrounded by large tracts of this invasive plant. It thus warrants further investigation whether chiggers not well fed by inferior *S. murinus* host in human residence area are required to be supplemented with chiggers well-fed by *R. losea* from surrounding *L. leucocephala* invasion sites, similar to the source-sink dynamics [76], and whether preserving *S. murinus*, one of the most abundant commensal mammals in Taiwan (Fig 8c; [77]), can help lower chigger population and thus human risks to scrub typhus.

As previous studies (e.g. [3, 7]), we investigated whether exotic plants sheltered more disease vectors than the other habitats. Unlike other studies, however, we expanded conventional spatial comparison to also track temporal vector population dynamics across seasons. We found that abundance of chiggers on the most important host, *R. losea*, was not only higher in invasion sites, seasonal fluctuation was also the lowest (with low CV), meaning that invasion sites have maintained a high and more stable chigger population. For example, during early winter (December), invasion sites still sustained a higher number of chiggers when the other two habitats sheltered few chiggers, suggesting that *L. leucocephala* could be a temporary refuge for chiggers and helped prolong chigger survival under less favorable climate. Likewise, abundance of ticks on *R. losea* was also higher and seasonally more stable in invasion sites. One exception is that abundance of ticks on *S. murinus* that hosted the highest proportion (around 70%) of ticks but not act as an ideal host (low engorgement), was also higher in invasion sites but displayed more dramatic seasonal fluctuation than human residence area. Such large fluctuation, however, was not due to low tick abundance in some months, but exceptionally high tick abundance in August.

Higher abundance of chiggers and ticks in *L. leucocephala* invasion sites than the other two habitats is less likely due to the habitat difference in microclimate because soil temperature and moisture was similar among the three habitats. Instead, the difference in chigger abundance might be attributed to invasion site’s harboring more *R. losea*, which was the ideal host of chiggers. Invasive plants have been shown to help aggregate and increase rodent abundance by providing dense cover from predators [78]. Likewise, *R. losea* might be sheltered under the dense *L. leucocephala* cover, which in turn increase the chigger abundance. On the other hand, unlike in chiggers that *R. losea* was evidently the most critical host, the relative contribution of *R. losea* and *S. murinus* to tick population was less clear. Relative to *R. losea* that most infested ticks were fully engorged, *S. murinus* was an inferior host but was infested with more ticks. Because there was more *S. murinus* surrounding human residence than outdoor fields (Fig 8c), as observed across Taiwan [77], higher abundance of ticks in invasion sites may thus be unrelated to the abundance of *S. murinus*. Instead, *R. losea* might be crucial in sustaining the high tick population. Similar to chiggers, elucidating the performance of ticks on *S. murinus* is pivotal for assessing whether *S. murinus* act as sinks for ticks and if preserving *S. murinus* can help lower the risks to tick-borne diseases.

Due to the inefficiency of sampling questing ticks and chiggers in Taiwan by such as flagging and black plate methods, vector abundance was not directly quantified, but instead by quantifying abundance of infested vectors on the trapped hosts. It might be argued that more vectors on the hosts would mean fewer vectors left on the ground questing for human victims. If this is true, however, hosts will soon be found lightly infested after the first cohort of emerged vectors finished their meal on the hosts and return to soils to molt or lay eggs. High abundance of chiggers and ticks on the hosts should thus reflect a continuous supply of questing chiggers and ticks in that habitat. The other limitation of this study is that in Penghu Islands, *L. leucocephala* is so exceedingly dominant that we were unable to include habitats inhabited solely by native plants for comparison. However, unlike native plants, eradicating *L. leucocephala* is extremely difficult due to its large soil seed bank. For example, in southern Taiwan the density can reach around 2,000 seeds per square meter [79]. Eradicating *L. leucocephala* for controlling vector-borne diseases will therefore be more challenging than removing native plants even when the latter habitat also shelters many disease vectors. From the public health perspective, investigating whether invasive sites are hotspots of vector-borne diseases thus warrant more concern.

This study highlights an important but largely neglected issue that a change in socioeconomy, such as a shift from agriculture to service and industry sectors, will stimulate a dramatic alteration on land use, which in turn can have considerable consequence for human risks to vector-borne diseases. The change in vegetative community accompanied with land use conversion, particular invasion of exotic plants, will not only interfere with the recovery of native fauna and flora but can potentially provide refuges for disease vectors and their vertebrate hosts under unfavorable weather, hence increasing disease burden for the general public. This is particularly worrying when eradicating invasive plants can often be very challenging.

## Author Contributions

CCK conceived and oversaw the research; CYW, JKW, HCS, HCW, CCK implemented the study, CCK did the statistical analyses; CYW, CCK wrote the original draft; CYW, JKW, HCS, HCW, CCK revised the final manuscript.

## Acknowledgements

We are indebted to members of Disease Ecology Lab. of National Taiwan Normal University for the help with the field work in Penghu. This study was financially supported by Taiwan Ministry of Science and Technology (MOST 104-2314-B-003-002-MY3; MOST 105-2621-M-003-003) awarded to CCK. The authors have no conflict of interest to declare.

## References

1. Kilpatrick AM, Randolph SE (2012) Drivers, dynamics, and control of emerging vector-borne zoonotic diseases. Lancet. 380:1946–1955.

2. Ogden NH, Lindsay LR (2016) Effects of climate and climate change on vectors and vector-borne diseases: ticks are different. Trends Parasitol 32:646–656.

3. Allan BF, Dutra HP, Goessling LS, Barnett K, Chase JM, et al. (2010) Invasive honeysuckle eradication reduces tick-borne disease risk by altering host dynamics. Proc Natl Acad Sci USA 107:18523–18527.

4. Lubelczyk CB, Elias SP, Rand PW, Holman MS, Lacombe EH, et al. (2004) Habitat Associations of *Ixodes scapularis* (Acari: Ixodidae) in Maine. Environ Entomol 33:900–906.

5. Elias SP, Lubelczyk CB, Rand PW, Lacombe EH, Holman MS, et al. (2006) Deer browse resistant exotic-invasive understory: an indicator of elevated human risk of exposure to *Ixodes scapularis* (Acari: Ixodidae) in southern coastal Maine woodlands. J Med Entomol 43:1142–1152.

6. Williams SC, Ward JS, Worthley TE, Stafford III KC (2009) Managing Japanese barberry (Ranunculales: Berberidaceae) infestations reduces blacklegged tick (Acari: Ixodidae) abundance and infection prevalence with *Borrelia burgdorferi* (Spirochaetales: Spirochaetaceae). Environ Entomol 38:977–984.

7. Williams SC, Ward JS (2010) Effects of Japanese barberry (Ranunculales: Berberidaceae) removal and resulting microclimatic changes on *Ixodes scapularis* (Acari: Ixodidae) abundances in Connecticut, USA. Environ Entomol 39:1911–1921.

8. Reiskind MH, Zarrabi AA (2011) The importance of an invasive tree fruit as a resource for mosquito larvae. J Vector Ecol 36:197–203.

9. Muturi EJ, Gardner AM, Bara JJ (2015) Impact of an alien invasive shrub on ecology of native and alien invasive mosquito species (Diptera: Culicidae). Environ Entomol 44:1308–1315.

10. Gardner AM, Muturi EJ, Overmier LD, Allan BF (2017) Large-scale removal of invasive honeysuckle decreases mosquito and avian host abundance. EcoHealth 14:750–761.

11. Gardner AM, Allan BF, Frisbie LA, Muturi EJ (2015) Asymmetric effects of native and exotic invasive shrubs on ecology of the West Nile virus vector *Culex pipiens* (Diptera: Culicidae). Parasit Vectors 8:329.

12. Civitello DJ, Flory SL, Clay K (2008) Exotic grass invasion reduces survival of *Amblyomma americanum* and *Dermacentor variabilis* ticks (Acari: Ixodidae). J Med Entomol 45:867–872.

13. Conley AK, Watling JI, Orrock JL (2011) Invasive plant alters ability to predict disease vector distribution. Ecol Appl 21:329–334.

14. Paris DH, Shelite TR, Day NP, Walker DH (2013) Unresolved problems related to scrub typhus: a seriously neglected life-threatening disease. Am J Trop Med Hyg 89:301–307.

15. Chikeka I, Dumler JS (2015) Neglected bacterial zoonoses. Clin Microbiol Infect 21:404–415.

16. Kelly DJ, Fuerst PA, Ching WM, Richards AL (2009) Scrub typhus: the geographic distribution of phenotypic and genotypic variants of *Orientia tsutsugamushi*. Clin Infect Dis 48 Supplement:S203–230.

17. Balcells ME, Rabagliati R, García P, Poggi H, Oddó D, et al. (2011) Endemic scrub typhus-like illness, Chile. Emerg Infect Dis 17:1659–1663.

18. Weitzel T, Dittrich S, López J, Phuklia W, Martinez-Valdebenito C, et al. (2016) Endemic scrub typhus in South America. N Engl J Med 375:954–961.

19. Kocher C, Jiang J, Morrison AC, Castillo R, Leguia M, et al. (2017) Serologic evidence of scrub typhus in the Peruvian Amazon. Emerg Infect Dis 23:1389–1391.

20. Thiga JW, Mutai BK, Eyako WK, Ng’ang’a Z, Jiang J, et al (2015) High seroprevalence of antibodies against spotted fever and scrub typhus bacteria in patients with febrile illness, Kenya. Emerg Infect Dis 21:688–691.

21. Horton KC, Jiang J, Maina A, Dueger E, Zayed A, et al. (2016) Evidence of *Rickettsia* and *Orientia* infections among abattoir workers in Djibouti. Am J Trop Med Hyg 95:462–465.

22. Maina AN, Farris CM, Odhiambo A, Jiang J, Laktabai J, et al. (2016) Q fever, scrub typhus, and rickettsial diseases in children, Kenya, 2011–2012. Emerg Infect Dis 22:883–886.

23. Roh JY, Song BG, Park WI, Shin EH, Park C, et al. (2014) Coincidence between geographical distribution of *Leptotrombidium scutellare* and scrub typhus incidence in South Korea. PLoS One 9:e113193.

24. Yang LP, Liang SY, Wang XJ, Li XJ, Wu YL, et al. (2015) Burden of disease measured by disability-adjusted life years and a disease forecasting time series model of scrub typhus in Laiwu, China. PLoS Negl Trop Dis 9:e3420.

25. Zheng L, Yang HL, Bi ZW, Kou ZQ, Zhang LY, et al. (2015) Epidemic characteristics and spatio-temporal patterns of scrub typhus during 2006-2013 in Tai’an, Northern China. Epidemiol Infect 143:2451–2458.

26. Cao M, Che L, Zhang J, Hu J, Srinivas S, et al. (2016) Determination of scrub typhus suggests a new epidemic focus in the Anhui Province of China. Sci Rep 6:20737.

27. Wu YC, Qian Q, Magalhaes RJS, Han ZH, Haque U, et al. (2016) Rapid increase in scrub typhus incidence in mainland China, 2006–2014. Am J Trop Med Hyg 94:532–536.

28. Harrison JL, Audy JR (1951) Hosts of the mite vector of scrub typhus II.-an analysis of the list of recorded hosts. Ann Trop Med Parasitol 45:186–194.

29. Traub R, Wisseman Jr. CL (1974) The ecology of chigger-borne rickettsiosis (scrub typhus). J Med Entomol 11:237–303.

30. Kawamura A, Tanaka H, Takamura A (1995) Tsutsugamushi Disease: An Overview. University of Tokyo Press, Tokyo.

31. Coleman RE, Monkanna T, Linthicum KJ, Strickman DA, Frances SP. et al. (2003) Occurrence of *Orientia tsutsugamushi* in small mammals from Thailand. Am J Trop Med Hyg 69:519–524.

32. Frances SP, Watcharapichat P, Phulsuksombati D (2001) Vertical transmission of *Orientia tsutsugamushi* in two lines of naturally infected *Leptotrombidium deliense* (Acari: Trombiculidae). J Med Entomol 38:17–21.

33. Phasomkusolsil S, Tanskul P, Ratanatham S, Watcharapichat P, Phulsuksombati D, et al. (2009) Transstadial and transovarial transmission of *Orientia tsutsugamushi* in *Leptotrombidium imphalum* and *Leptotrombidium chiangraiensis* (Acari: Trombiculidae). J Med Entomol 46:1442–1445.

34. Shatrov AB (2000) On the origin of parasitism in trombiculid mites (Acariformes: Trombiculidae). Acarologia 41:205–213.

35. Parola P, Paddock CD, Raoult D (2005) Tick-borne rickettsioses around the world: emerging diseases challenging old concepts. Clin Microbiol Rev 18:719–756.

36. Parola P, Paddock CD, Socolovschi C, Labruna MB, Mediannikov O, et al. (2013) Update on tick-borne rickettsioses around the world: a geographic approach. Clin Microbiol Rev 26:657–702.

37. Raoult D, Roux V (1997) Rickettsioses as paradigms of new or emerging infectious diseases. Clin Microbiol Rev 10:694–719.

38. Needham GR, Teel PD (1991) Off-host physiological ecology of ixodid ticks. Annu Rev Entomol 36:659–681.

39. Stafford KC (1994) Survival of immature *Ixodes scapularis* (Acari, Ixodidae) at different relative humidities. J Med Entomol 31:310–314.

40. Randolph SE (2004) Tick ecology: processes and patterns behind the epidemiological risk posed by ixodid ticks as vectors. Parasitology 129:S37–S65.

41. Foley JA, DeFries R, Asner GP, Barford C, Bonan G, et al. (2005) Global consequences of land use. Science 309:570–574.

42. Cramer VA, Hobbs RJ, Standish RJ (2008) What’s new about old fields? Land abandonment and ecosystem assembly. Trends Ecol Evol 23:104–112.

43. Aide TM, Grau HR (2004) Globalization, migration, and Latin American ecosystems. Science 305:1915–1916.

44. Rudel TK, Coomes OT, Moran E, Achard F, Angelsen A, et al. (2005) Forest transitions: towards a global understanding of land use change. Global Environ Chang 15: 23–31.

45. Grau HR, Aide M (2008) Globalization and land-use transitions in Latin America. Ecol Soc 13:16.

46. Lasanta T, Arnáez J, Pascual N, Ruiz-Flaño P, Errea MP, et al. (2017) Space–time process and drivers of land abandonment in Europe. Catena 149:810–823.

47. Benayas JMR, Martins A, Nicolau JM, Schulz JJ (2007) Abandonment of agricultural land: an overview of drivers and consequences. CAB Reviews: Perspectives in Agriculture, Veterinary Science, Nutrition and Natural Resources 2, No.057.

48. Grau HR, Aide TM, Zimmerman JK, Thomlinson JR, Helmer E, et al. (2003) The ecological consequences of socioeconomic and land-use changes in postagriculture Puerto Rico. BioScience 53:1159–1168.

49. Parés-Ramos I, Gould W, Aide T (2008) Agricultural Abandonment, Suburban Growth, and Forest Expansion in Puerto Rico between 1991 and 2000. Ecol Soc 13:1.

50. Lugo AE, Helmer E (2004) Emerging forests on abandoned land: Puerto Rico’s new forests. Forest Ecol Manag 190:145–161.

51. Hsu HC (2005) A Further Documentary of Penghu County. Vol. 5. Natural Products. Penghu County Government. (In Chinese)

52. Urban and Regional Development Statistics 2017. National Development Council, Taiwan. (In Chinese)

53. Agricultural Statistics Yearbook 2015. Council of Agriculture, Executive Yuan, Taiwan.

54. Lowe S, Browne M, Boudjelas S, De Poorter M (2000) 100 of the World’s Worst Invasive Alien Species A selection from the Global Invasive Species Database. Published by The Invasive Species Specialist Group (ISSG) a specialist group of the Species Survival Commission (SSC) of the World Conservation Union (IUCN), 12pp.

55. Huang CY (2009) Distribution, germination characters, allelopathy and chemical control of *Leucaena leucocephala* (Lam.) de Wit in Penghu. Master Thesis. Department of Agronomy, National Chung Hsing University. (In Chinese, English abstract)

56. Cooper WC, Lien JC, Hsu SH, Chen WF (1964) Scrub typhus in the Pescadores Islands: an epidemiologic and clinical study. Am J Trop Med Hyg 13:833–838.

57. Lien JC, Liu SY, Lin HM (1967) Field observation on the bionomics of *Leptotrombidium deliensis*, the vector of scrub typhus in the Pescadores. Acta Medica et Biologica 15(Suppl):27–31.

58. Dirk Van Peenen PF, Lien JC, Santana FJ, See R (1976) Correlation of chigger abundance with temperature at a hyperendemic focus of scrub typhus. J Parasitol 62:653–654.

59. Olson JG (1979) Forecasting the onset of a scrub typhus epidemic in the Pescadores Islands of Taiwan using daily maximum temperatures. Trop Geogr Med 31:519–524.

60. Olson JG, Scheer EJ (1978) Correlation of scrub typhus incidence with temperature in the Pescadores Island of Taiwan. Ann Trop Med Parasitol 72:195–196.

61. Olson JG, Ho CM, Van Peenen PFD, Santana FJ (1978) Isolation of *Rickettsia tsutsugamushi* from mammals and chiggers (Fam. Trombiculidae) in the Pescadores Islands, Taiwan. Trans R Soc Trop Med Hyg 72:192–194.

62. Olson JG, Bourgeois AL, Fang RC (1982) Population indices of chiggers (*Leptotrombidium deliense*) and incidence of scrub typhus in Chinese military personnel, Pescadores Islands of Taiwan, 1976-77. Trans R Soc Trop Med Hyg 76:85–88.

63. Kuo CC, Shu PY, Mu JJ, Wang HC (2015) High prevalence of *Rickettsia* spp. infections in small mammals in Taiwan. Vector Borne Zoonotic Dis 15:13–20.

64. Kuo CC, Shu PY, Mu JJ, Lee PL, Wu YW, et al. (2015) Widespread *Rickettsia* spp. infections in ticks in Taiwan. J Med Entomol 52:1096–1102.

65. Keesing F, Brunner J, Duerr S, Killilea M, LoGiudice K, et al. (2009) Hosts as ecological traps for the vector of Lyme disease. Proc R Soc Lond B Biol Sci 276:3911–3919.

66. Kuo CC, Wang HC, Huang CL (2011) Variation within and among host species in engorgement of larval trombiculid mites. Parasitology 138:344–353.

67. Li J, Wang D, Chen X (1997) Trombiculid Mites of China: Studies on Vector and Pathogen of Tsutsugamushi Disease. Guangdong Science & Technology Publishing, Guangzhou. (In Chinese)

68. Teng KF, Jiang ZJ (1991) Economic Insect Fauna of China. Fasc 39. Acari: Ixodidae. Editorial Committee of Fauna Sinica, Academic Sinica. Science Press, Beijing. (In Chinese)

69. Black WC, Piesman J (1994) Phylogeny of hard- and soft-tick taxa (Acari: Ixodida) based on mitochondrial 16S rDNA sequences. Proc Natl Acad Sci USA 91:10034–10038.

70. Beati L, Keirans JE (2001) Analysis of the systematic relationships among ticks of the genera *Rhipicephalus* and *Boophilus* (Acari: Ixodidae) based on mitochondrial 12S ribosomal DNA gene sequences and morphological characters. J Parasitol 87:32–48.

71. Kuo CC, Huang JL, Shu PY, Lee PL, Kelt DA, et al. (2012). Cascading effect of economic globalization on human risks of scrub typhus and tick-borne rickettsial diseases. Ecol Appl 22:1803–1816.

72. Wood M (2005) Bootstrapped confidence intervals as an approach to statistical inference. Organ Res Methods 8:454–470.

73. Heinze G (2006) A comparative investigation of methods for logistic regression with separated or nearly separated data. Stat Med 25:4216–4226.

74. Kuo CC, Lee PL, Chen CH, Wang HC (2015) Surveillance of potential hosts and vectors of scrub typhus in Taiwan. Parasit Vectors 8:611.

75. LoGiudice K, Ostfeld RS, Schmidt KA, Keesing F (2003) The ecology of infectious disease: effects of host diversity and community composition on Lyme disease risk. Proc Natl Acad Sci USA 100:567–571.

76. Pulliam HR (1988) Sources, sinks, and population regulation. Am Nat 132:652–661.

77. Chang CH, Lin JY, Lin LK, Yu JYL (1999) Annual reproductive patterns of female house shrew, *Suncus murinus*, in Taiwan. Zool Sci 16:819–826.

78. Malo AF, Godsall B, Prebble C, Grange Z, McCandless S, et al. (2012) Positive effects of an invasive shrub on aggregation and abundance of a native small rodent. Behav Ecol 24:759–767.

79. Lin CC (2011) Characteristics of seed germination and seedling regeneration of *Leucaena leucocephala*. Master Thesis. Department of Forest, National Pingtung University of Science and Technology. (In Chinese, English Abstract)

